# Laser-free super-resolution microscopy

**DOI:** 10.1101/121061

**Authors:** Kirti Prakash

## Abstract

We report that high-density single-molecule super-resolution microscopy can be achieved with a conventional epifluorescence microscope setup and a Mercury arc lamp. The configuration termed as laser-free super-resolution microscopy (LFSM), is an extension of single molecule localisation microscopy (SMLM) techniques and allows single molecules to be switched on and off (a phenomenon termed as “blinking”), detected and localised. The use of a short burst of deep blue excitation (350-380 nm) can be further used to reactivate the blinking, once the blinking process has slowed or stopped. A resolution of 90 nm is achieved on test specimens (mouse and amphibian meiotic chromosomes). Finally, we demonstrate that STED and LFSM can be performed on the same biological sample using a simple commercial mounting medium. It is hoped that this type of correlative imaging will provide a basis for a further enhanced resolution.

## Introduction

The resolution of images generated by light microscopy is limited by diffraction due to the wave nature of light (Born and Wolf, 1980; Betzig and Trautman, 1992; Cremer and Masters, 2013). As a consequence, in the image plane, a pointlike object is registered as a blurred Airy-disc. The function that describes this blurring is referred to as the Point Spread Function (PSF) and can be captured in terms of the numerical aperture (NA) of the objective lens and the wavelength (*λ*) of the light used. When the most optimal combination of objective lens and light in the visible region is used, a lateral resolution of about 200 nm can be achieved, which is often referred to the resolution limit of light microscopes (Abbe, 1873).

The resolution limit of light microscopes has been recently overcome by a number of superresolution microscopy techniques (SMTs), such as single molecule localization microscopy (SMLM) (Lidke et al., 2005; Betzig et al., 2006; Rust et al., 2006; Hess et al., 2006), stimulated emission depletion (STED) (Hell and Wichmann, 1994; Willig et al., 2006), structured illumination microscopy (SIM) (Heintzmann and Cremer, 1999; Gustafsson, 2000) and various related techniques (Schwentker et al., 2007; Heilemann et al., 2008; Schoen et al., 2011; Szczurek et al., 2014; Dertinger et al., 2009; Gustafsson et al., 2016; Martens et al., 2019).

All current superresolution techniques make use of coherent light sources such as lasers, which in most cases are a necessity, for instance, in STED. For single molecule based superresolution techniques a minimum power (~0.1 kW/*cm*^2^) is required to switch the fluorophores between dark and bright state (Dickson et al., 1997; Betzig et al., 2006). For example, ~10 mW on 100 × 100 *μm^2^* in the sample plane would typically be enough for exciting and detecting single molecules. Another wavelength light (for example, 405 nm laser) can then be used to trigger switching from the dark to the bright state.

We wondered if one could make use of an incoherent light source instead of a laser to induce the on/off switching of single molecules, since previously, it has been shown that Mer-cury arc lamps and LEDs can be used for detection of single molecules (Chiu and Quake, 1999; Gerhardt et al., 2011). For reactivation, we found 350-380 nm peaks (instead of a 405 nm laser) of the Mercury lamp can be used to trigger switching from the dark to the bright state. Thus, a Hg lamp overcomes the need of any lasers and moving the single-molecule field towards laser-free super-resolution microscopy (LFSM). In this report, we demonstrate that using a Mercury arc lamp and appropriate combination of dyes, imaging buffer and filters, one can extract enough photons to detect, localise and reactivate a fluorophore with a nanometer precision.

#### Plain English

Fifteen years into its development, super-resolution microscopy is still limited to relatively few microscopy and optics groups. This is mainly due to the significant cost of current superresolution microscopes, which require high-quality lasers, high NA objective lenses, very sensitive cameras, and highly precise microscope stages, and to the complexity of post-acquisition data reconstruction and analysis. We present results that demonstrate the possibility of obtaining nanoscale-resolution images using a conventional microscope and an incoherent light source. We describe an easy-to-follow protocol that every biologist can implement in the laboratory. We hope that this finding will help any scientist to generate high-density superresolution images even with a limited budget. Ultimately, the new photophysical observations reported here should pave the way for more in-depth investigations on the processes underlying the excitation, photobleaching and photoactivation of a fluorophore.

We applied a simple protocol on a standard epifluorescence microscope setup to generate high-resolution images of synaptonemal complexes (SCs) and lampbrush chromosomes (LBCs) and compared the results with those produced by state-of-the-art superresolution techniques. In our configuration, single fluorescent molecules undergo repeated cycles of fluorescent bursts, similarly to what can be observed with other SMLM approaches. The individual blinking events are detected with a standard CCD camera and localised with high precision. The positions of individual molecules are then used to reconstruct an image with high spatial resolution. The novel features of our setup are the following:

- On/off switching of single molecules by a Mercury arc lamp instead of coherent light sources such as lasers.
- High-resolution single-molecule images with an unmodified epifluorescence microscope, as commonly found in cell biology laboratories everywhere, instead of a specially constructed instrument.
- The use of a short burst of deep blue excitation (350-380 nm, Mercury arc lamp with a DAPI filter, 365/30 nm) for a prolonged reactivation of molecules, once blinking has slowed or stopped. Previously, either a 405 nm (Mercury arc lamp with a line filter) or a 405 nm laser was used for photoswitching (Dickson et al., 1997; Betzig et al., 2006). The prolongated blinking helped to reconstruct super-resolved images with a high signal density.
- Ability to perform STED and SMLM measurements on the same biological sample employing a simple imaging medium (ProLong Diamond).

Regarding the different observations listed above, and with the growing interest of the community for validation of experiments at the nanoscale level, we believe that the results presented in this paper potentially have a very broad application.

## Results

### Description of the LFSM setup

Mercury arc lamp is a common source of illumination in most epifluorescence microscopes (Figure 1A). Such lamps emit light with peaks around 400 nm and 560 nm (Figure 1B, S3). Several dyes have been designed so that their absorption spectra corresponds to thes peaks of the Mercury arc lamp for optimal fluorescence. One such dye, Texas Red, has its excitation peak around 594 nm. In our experiments, we used Alexa Fluor 594, an alternative version of Texas Red, for its brighter signal and photo-stability.

**Figure 1.**
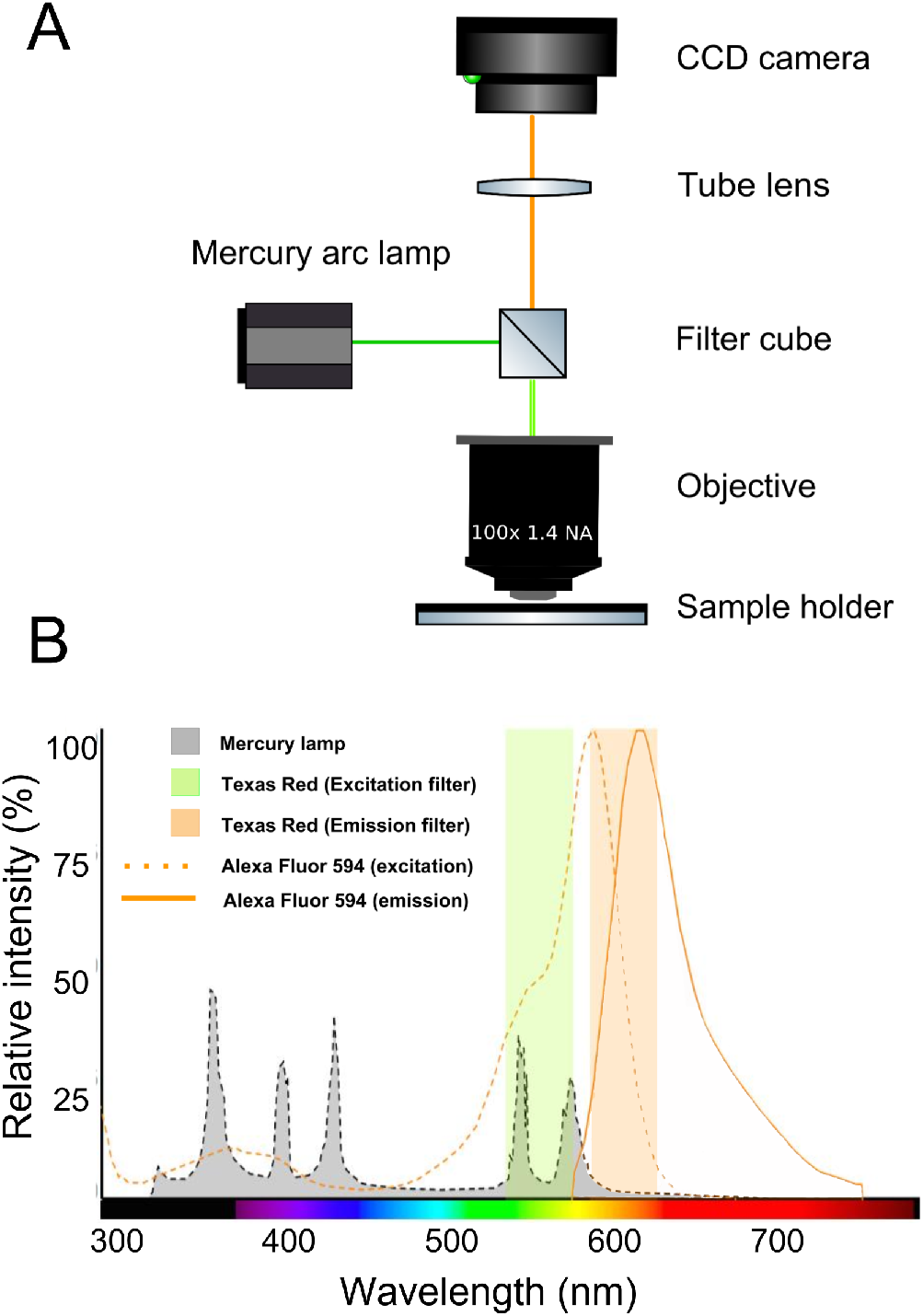
Laser-free super-resolution microscopy (LFSM) setup: (A) Schematic of the optical system of a standard epifluorescence microscope, as used in LFSM. (B) The shaded grey area shows the emission spectrum of a Mercury arc lamp. We use the peaks around 540-580 nm to excite the fluorophore (Alexa 594) and the peaks around 350-380 nm to reactivate the fluorophore. The fluorophore then begins to blink, emitting light with a maximum around 620 nm.

For our measurements, we used Olympus BX61 (Figure S1), a standard widefield setup. The microscope was equipped with a Mercury arc lamp, a 100x objective lens (1.40 N.A.) and a digital CCD camera. In the illumination module, we used a Texas Red excitation filter (560/40 nm), which transmits all wavelengths between 540 and 580 nm peaks of the Mercury arc lamp (Figure 1B).

We start to record images once the individual molecules began to blink after the initial bleaching step (approximately 10-20 minutes), but the time depends on the light intensity, dye density and imaging medium. For optimizing imaging conditions for single-molecule localization imaging, please refer (Diekmann et al., 2020). After 1-2 min of imaging with Texas Red filter, the blinking diminishes and it is necessary to switch to the DAPI filter (365/30 nm, 30 seconds) to reactivate the molecules. The process of photobleaching and subsequent photoactivation for a sparse set of molecules needs to be repeated (approximately 1 to 2 hours) until sufficient signal density is reached for a high-density image reconstruction.

On the microscopy side, use of the Mercury arc lamp helps to avoid the problems usually associated with lasers: alignment, the need for multiple spectral lines, the expense, danger to the eyes and the premature bleaching of the fluorophores. However, the main drawback using a Mercury arc lamp is that one cannot reactivate at the same time as imaging. Finally, the use of ProLong Diamond as the imaging medium further minimised the effort to prepare a complex cocktail of reducing/oxidising reagents to create a redox environment (Heilemann et al., 2008; Löschberger et al., 2012).

### On/off switching of single molecules induced by LFSM

To test whether individual molecules can be switched on/off with an incoherent light source, we labelled gp210, a protein known to occupy the nuclear pore complex periphery, with Alexa Fluor 594. ProLong Diamond was as the anti-fading mounting solution. The composition of ProLong Diamond is unknown and might not provide the classic redox environment as needed for Alexa Fluor 647. However, simple buffers such as Vectashield have been previously used for single-molecule imaging of Alexa Fluor 647 (Olivier et al., 2013).

Initially, the dye fluoresces as usual. Over a matter of minutes the sample gradually bleaches entirely, but then individual fluorophore begin to ‘blink.’ That is, they fluoresce for a short time, then cease to fluoresce, and then fluoresce again. The blinking events are recorded with a standard CCD camera (Betzig et al., 2006).

To quantify the switching of single molecules, we chose an area of 7X7 pixels (roughly 450X450 *nm*^2^) and plotted the intensity profile along the image stack (Figure 2A-C). We could observe sharp peaks, which represent fluorescent bursts of a single molecule integrated over 150 ms of camera exposure. The broad peaks in the profile occur when a fluorophore remains ‘on’ for an extended period or when the neighboring fluorophores within the same diffraction limited spot fluoresce during the same time span. As we have no way to isolate such fluorophores optically, we merged all such signals in the consecutive frames during the final analysis (see Methods and Materials section for more details).

**Figure 2.**
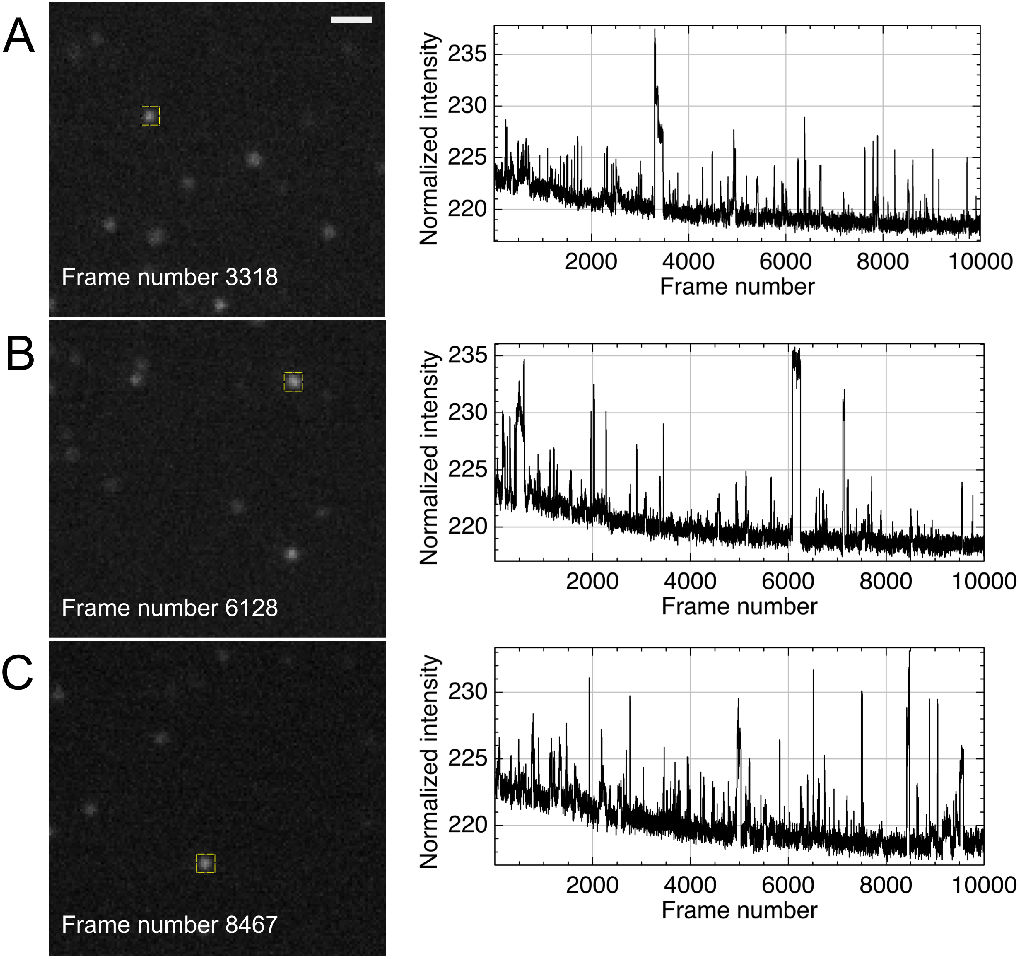
Proof of on/off switching of single-molecules induced by an incoherent light source: Blinking of single molecules (gp210, a nuclear pore complex protein labelled with Alexa Fluor 594) is shown using a Mercury arc lamp. (A-C) Three selected frames (3318, 6128 and 8467) out of 10000 frames are shown on the left side where a subset of molecules are ‘on’. On the right side, intensity profiles for three 7X7 pixels (roughly 450X450 *nm^2^*) cross sections (yellow boxes in the images on the left) along the image stack are shown. Fluorescent bursts of single molecules can be observed as sharp peaks. Scale bar: 1 *μm* in A and is same for B, C.

### Superresolution microscopy with LFSM

Next, we wondered if the positions of single molecule could be used to reconstruct a super-resolved image. We immunostained SYCP3 (SYnaptonemal Complex Protein 3) with Alexa Fluor 594. SYCP3 forms the lateral component of SC and is an ideal structure for benchmarking of superresolution microscopes as these lateral elements are 150 nm apart and cannot be resolved using a conventional light microscope (Figure 3A-B). Using LFSM, we generated localisation maps of SYCP3 by integrating approximately 50000 observations, each of which captured photons emitted during 150 ms of camera integration time, and were now able to resolve the two strands of the SYCP3 (Figure 3C). A more precise comparison between confocal and LFSM images is quantified using line scans (Figure 3D). On average, we detected 1000 photons per signal (Figure 3E) with a localization precision of 20 nm (Figure 3F). We achieved a Fourier Ring Correlation (FRC) resolution (Nieuwenhuizen et al., 2013) of about 90 nm (Figure 3G) in the case of SC samples.

**Figure 3.**
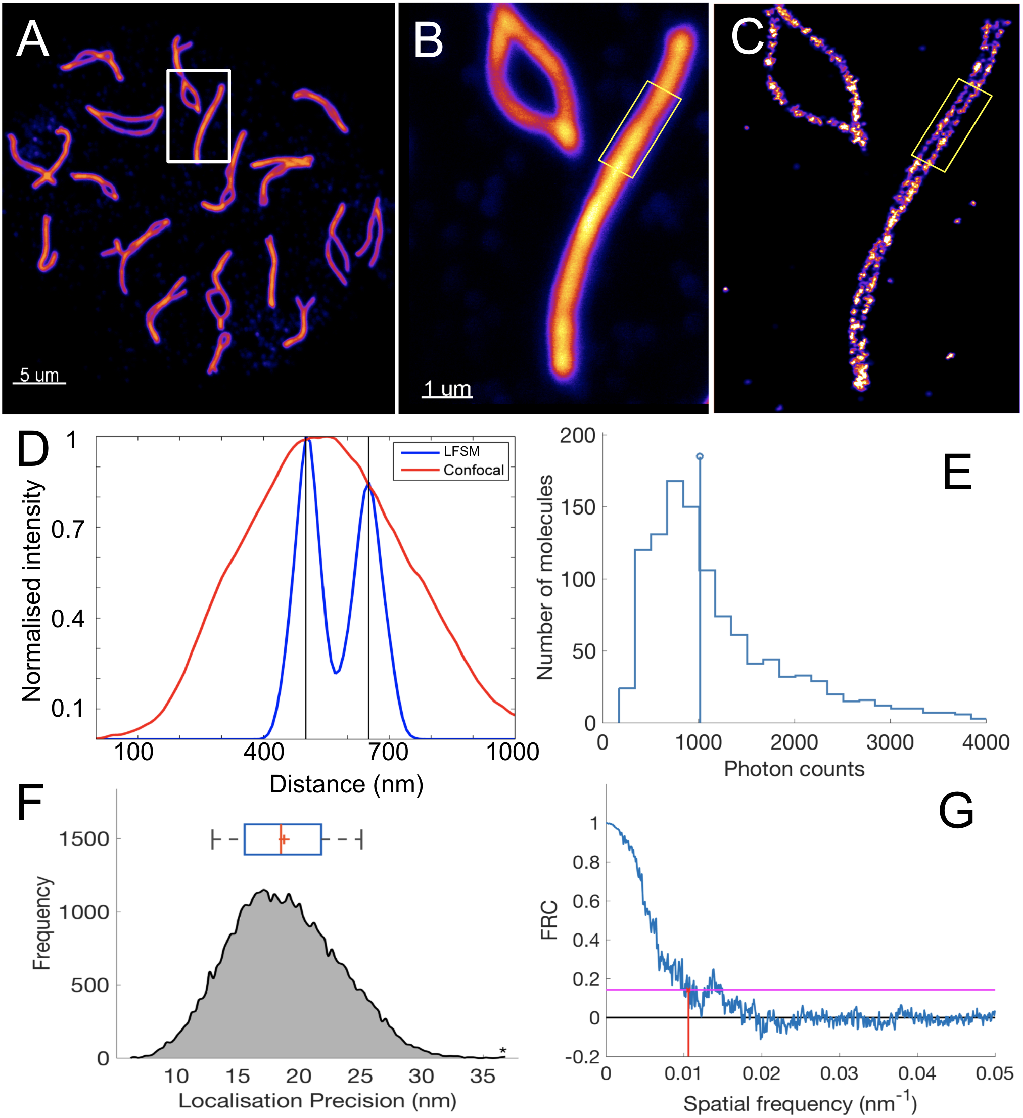
Single molecule superresolution microscopy with LFSM: Pachytene chromosomes from mouse stained with an antibody against the synaptonemal complex (SC). (A) The low magnification confocal image shows the entire complement of chromosomes. (B) The boxed area in (A) shown at a higher magnification of the confocal microscope. Note that the two halves of the SC that unite the meiotic chromosome pair are not resolved. (C) The same chromosome imaged with LFSM and the two halves of the SC could be resolved. (D) The contrast between LFSM and confocal microscopy is further demonstrated by line scans of the boxed regions in B and C. Line scans show that LFSM can resolve the two halves of the synaptonemal complex (about 150 nm separation). (E) An average number of photons (median: 1000 photons/signal) extracted from single molecules after illumination with a Mercury arc lamp. (F) An average localisation precision of 20 nm is obtained for Alexa Fluor 594. The blue box above the plot denotes the 25% and 75% quartiles while the whiskers bound 9% and 91% of the data. The red line in the box denotes the median while the ‘+’ sign indicates the mean. (G) FRC resolution around 90 nm is currently being achieved with LFSM for SC samples. The inverse of spatial frequency (red line) provides an estimate for the resolution. The horizontal pink line indicates the 1/7 threshold of the radially combined Fourier frequencies as suggested by (Nieuwenhuizen et al., 2013). Scale bar: 5 *μ*m in A. 1 *μ*m in B and is same for C.

We further compared localisation maps of SYCP3 with LFSM and that of a standard high-end SMLM setup (Figure S2A-B), each of which captured photons emitted during 150 ms and 100 ms of camera integration time, respectively. SMLM data taken from (Prakash et al., 2015) for comparison with LFSM. On average, we detected 1000 photons per cycle, which is comparable to the number of photons per cycle we get with Alexa Fluor 555 using a standard SMLM (Figure S2C). The setups localised individual fluorophores with an average precision of 11 nm (standard SMLM) and 18 nm (OLM) (Figure S2D). Presently, we achieve a FRC resolution of 67 nm for Alexa Fluor 555 when illuminated with a laser (Figure S2E), and a FRC resolution of 94 nm for Alexa Fluor 594 when illuminated with a Mercury arc lamp (Figure S2F).

### Photo reactivation of single molecules provides high signal density

We next discovered that 350-380 nm spectral peaks in the Mercury arc lamp could also be used for the photoactivation of single molecules and can make a fluorochrome blink for an extended recording of the data. A continuous exposure of the sample with the Mercury arc lamp (tested up to 12 hours) did not lead to permanent bleaching (Figure 4A-D). We define permanent bleaching when the blinking process of the fluorophores cannot be reactivated even after UV illumination. One of the current problems with localisation microscopy is that the high laser power tends to bleach the sample before one reaches a signal density sufficient to make biological inferences. This reactivation will be useful for experiments where the underlying structure is unknown and a lot of localisations are needed to bring out the structure. Moreover, it might be possible that the relatively low power and non-coherent nature of the Mercury arc lamp helped to prevent the permanent bleaching of the sample (Diekmann et al., 2020).

**Figure 4.**
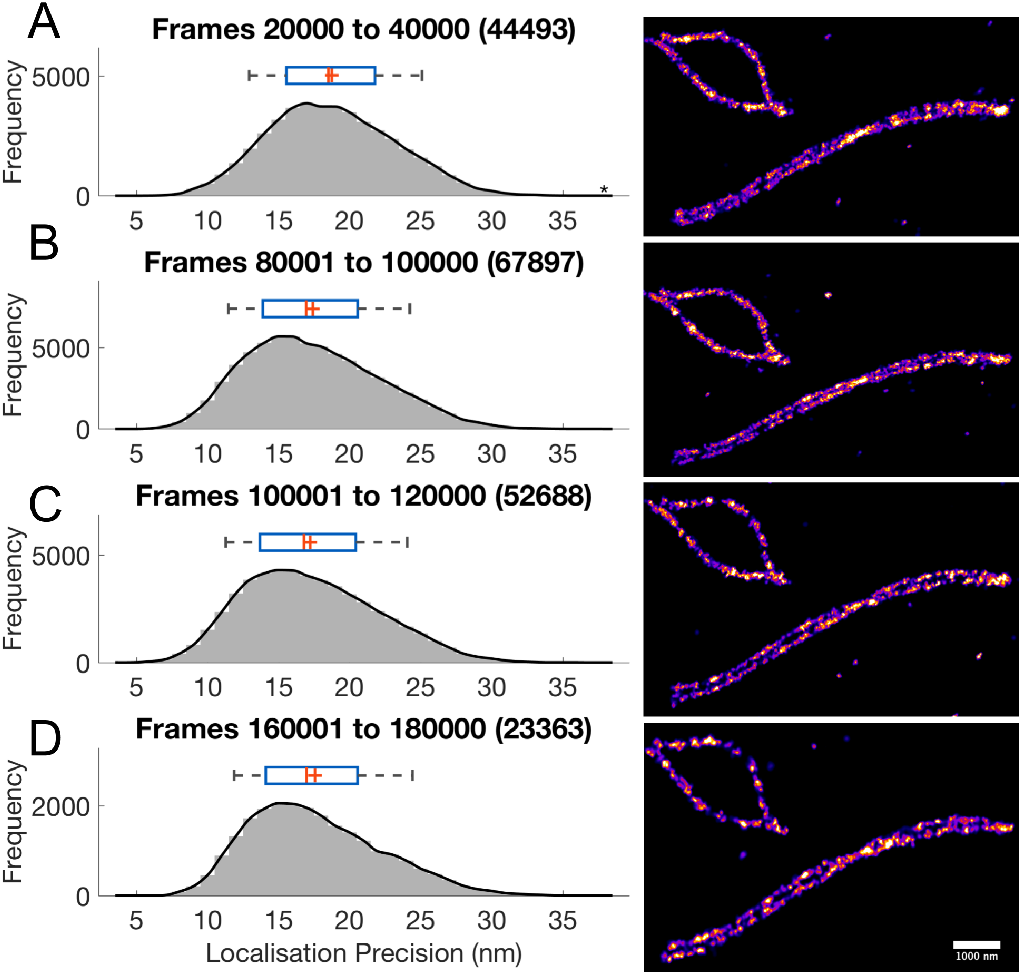
Photo reactivation of single molecules and prolonged switching of single molecules with the Mercury arc lamp. (A-D) shows independent acquisitions of SC labelled with Alexa Fluor 594. The mean localisation precision (approx. 20 nm) was consistent over the extended period of data acquisition. (A) In total, 44493 signals were acquired in a period over the course of 20000 frames. (B) Next 80000-100000 frames with 67897 signals, (C) The subsequent 20000 frames with 52688 signals and (D) The final 20000 frames with 23363 signals. Signals were recorded over a period of 12 hours. The blue box above each plot denotes the 25% and 75% quartiles while the whiskers bound 9% and 91% of the data. The red line in the box for each plot denotes the median while the ‘+’ is for the mean. Scale bar: 1 *μm* in D and is same for A, B, C.

We would like to stress here that high signal density with good localisation accuracy and the low duty cycle are critical to reconstruct high-resolution images from the single molecule data (Dempsey et al., 2011; Legant et al., 2016; Ha and Tinnefeld, 2012). With indefinite blinking of Alexa Fluor 594, obtaining high number of signals with a good precision is easy, however, it has to be complemented with a low duty cycle and a higher localisation accuracy of the fluorescent moiety.

### Photo reactivation can be used for correlative LFSM and STED

Next, we observed that using a DAPI excitation filter (365/30 nm), we could recover the sample after STED bleaching. Figure 5A-B shows a bleached cross section of the nuclear envelope after STED acquisition. We used spectral peaks (range 350-380 nm) of the Mercury arc lamp to recover the bleached sample (Figure 5C).

**Figure 5.**
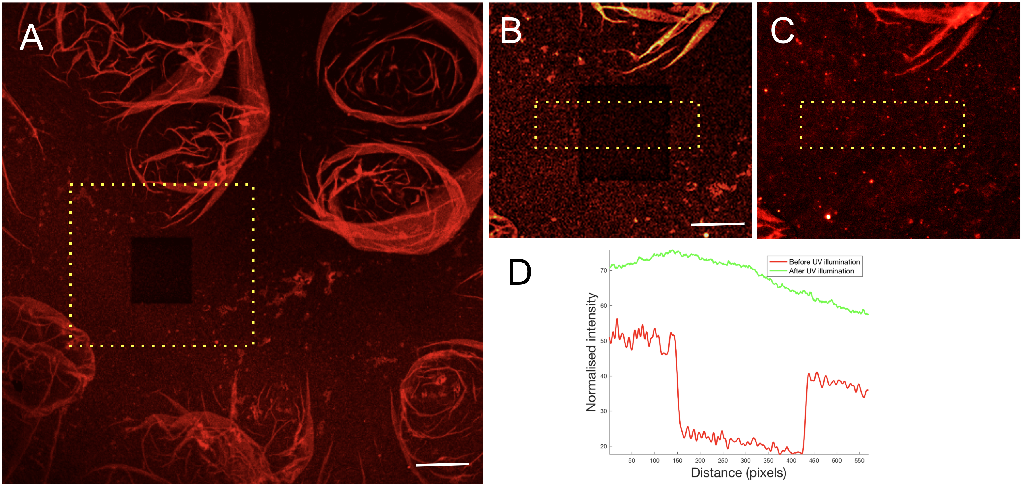
Photo reactivation of a nuclear membrane (stained with Alexa 594) cross section using the Mercury arc lamp: (A) shows the bleached cross-section after STED imaging. (B) The section in the yellow box in (A). (C) The same photobleached section as in (B) after the photoactivation. (D) Normalized intensity comparison (yellow box in B and C) before and after UV illumination. Scale bar: 2.5 *μ*m in A. 1 *μ*m in B and is same for C.

Alexa Fluor 594, due to its brightness and photostability, also happens to be a good STED fluorochrome. Moreover, the STED depletion laser wavelength does not excite the dye. We wondered if we could make use of the reactivation property of Alexa Fluor 594 with UV illumination to do correlative LFSM and STED. We first imaged Alexa Fluor 594 in the confocal mode (Figure 6A) with Leica SP8 and then in STED mode (Figure 6D), with 660 and 775 nm depletion lasers. Next, we imaged the same region of the sample on an Olympus BX 61 in the widefield mode (Figure 6B). Once bleached, we used UV illumination (365/30 nm filter) to recover the signal and then Texas Red filter to excite a subset of photo-activated molecules for LFSM imaging (Figure 6E). The minor structural differences in LFSM and STED images are partially due to the thickness of the lampbrush chromosomes (approximately 1 *μ*m) and partially due to imaging of slightly different sample planes.

**Figure 6.**
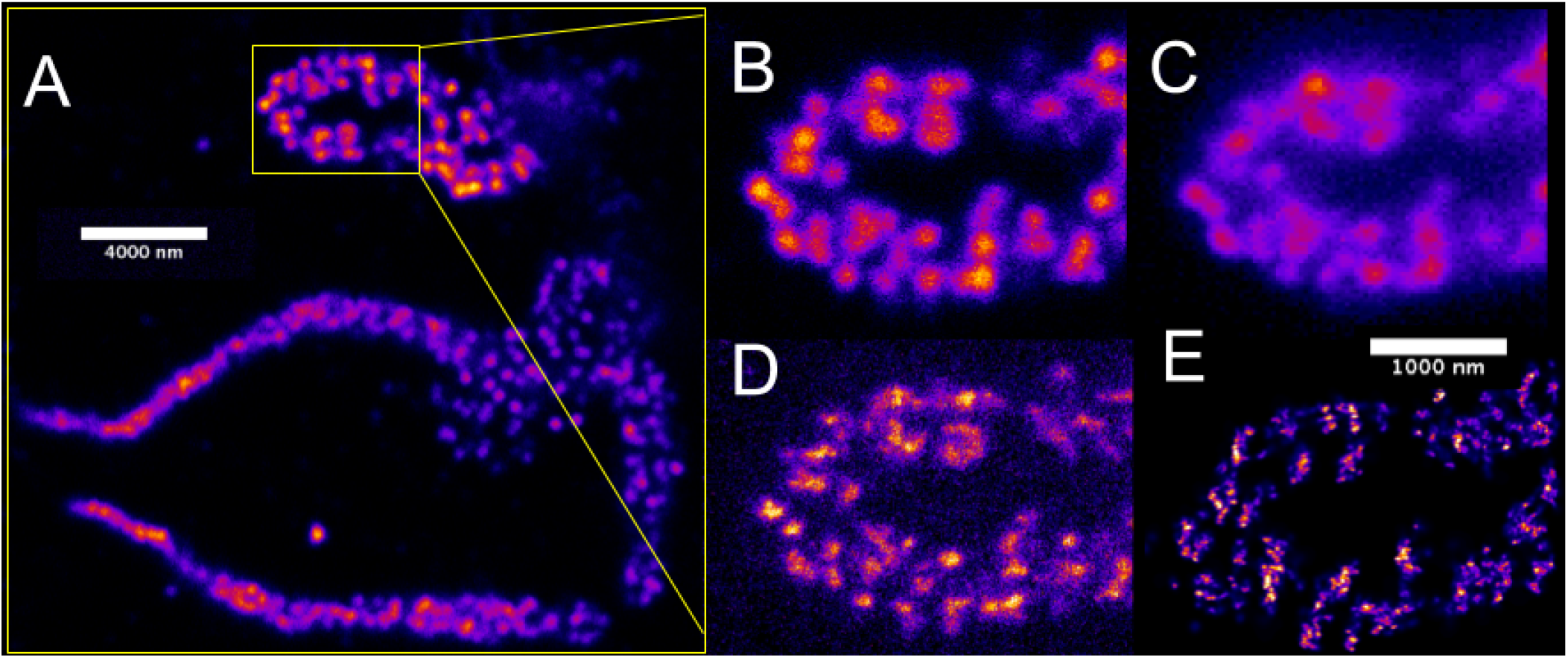
LFSM and STED microscopy: The distribution of the RNA-binding protein CELF1 stained with Alexa Fluor 594 on specific loops of lampbrush chromosomes (LBCs) is compared using different light microscopy techniques. (A) The low magnification confocal image shows the entire complement of the lampbrush chromosomes. (B) The boxed area in A shown at a higher magnification using confocal (B), widefield (C), STED (D) and LFSM (E). Scale bar: 4 *μ*m in A. 1 *μ*m in E and is same for B, C, D.

## Methods

All experimental procedures were performed in compliance with ethical regulations and approved by the IACUC of the Carnegie Institution for Science. The meiotic spreads in mouse oocyte, nuclear pore complex samples, and the lampbrush chromosome samples were kindly provided by Joseph G Gall, Zehra Nizami, and Safia Malki. Details regarding the preparation of samples can be found in the following articles (Susiarjo et al., 2009; Malki and Bortvin, 2017; Prakash et al., 2015; Gall and Nizami, 2016; Shi et al., 2017).

### Imaging medium

We rinsed coverslips in water and used ProLong Diamond (ThermoFIsher, P36970) as the anti-fading mounting solution. The refractive index of the oil used was 1.518. We wish to emphasise here that ProLong Diamond works successfully for both STED and LFSM/SMLM microscopy with Alexa Fluor 594. For two or more colours imaging, a more complex and optimised imaging buffer might be required.

### Microscopy

Confocal and STED images were obtained using a Leica TCS SP8 microscope with 592-nm, 660-nm, and 775-nm depletion lasers. LFSM images were acquired with an Olympus BX61 microscope equipped with 100x / NA 1.4 oil objective lens (Olympus UPlanSApo) and a Hamamatsu CCD camera (C4742-95). The effective pixel size was 64.5 nm in the sample plane. Illumination was done using a Mercury arc lamp (100W, Ushio USH-103D), as is commonly used in the conventional epifluorescence microscopes. We detected 50 mW power in the sample plane with a circular detector (10 mm in diameter, X-Cite XR2100). The filter cube consisted of the following excitation filters: Texas Red (560/40 nm) and DAPI (365/30), emission filter (590/40 nm) and a dichromatic mirror. All the filters were bought from Semrock. The microscope was placed on a simple table with no stabilisation or control for vibration (Figure S1). Due to this, there was a considerable drift of the sample during the measurements. Due to reactivation of Alexa 594 with DAPI excitation filter, we could record a high number of frames, which provided us with the luxury to discard the frames with significant drift. The frames with less drift were corrected using post-acquisition drift correction algorithms as described in (Prakash, 2017, 2016).

### Data acquisition

A few parameters such as bleaching time, the number of signals per frame, camera integration time and the total number of final frames need to be optimised before acquiring the data. In our case, 10-20 minutes of pre-bleaching with Texas Red excitation filter is required before molecules start to blink. The pre-bleaching step further helps to minimise autofluorescence which was minimal in the case of thin samples like synaptonemal complex (SC). Once blinking started, we adjusted the lamp power to have 10-20 signals per frame in a cross section of 25 μm^2^. Only a few signals per frame help to optically isolate the molecules from each other and obtain high-resolution images. We found approximately 20000 frames with 50000 localizations to be enough to reconstruct high-density images of SC. An integration time of 100-150 ms was sufficient for a good signal-to-noise ratio. Finally, to speed up the acquisition time and save data storage space, the imaging area was restricted to the region of interest.

### Data reconstruction and visualisation

After data acquisition, the coordinates of the single signals need to be precisely determined. Figure S4 presents an overview of various steps required to reconstruct a highly resolved image from a stack comprising a large number of images, where each image contains only a few signals.

As the first step, single molecule signals in each frame need to be separated from the background. The background varies throughout the image stacks, but the variation in the background between the consecutive images is relatively negligible. We used this strategy to estimate the background for a given frame based on the information from the previous images. The noise in the image can further make it difficult to determine the coordinates of the signal precisely. The error in the measured intensity *N* is the square root of the measured value 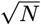, following a Poisson distribution. This error can become significant when the background signal is high and non-uniform. In the present case, we estimated a background map for each image by averaging previous ten frames and then subtracting from the image to get the difference image. Next, a median filter was applied to the difference image to get the signals above an empirically determined threshold.

In the second step, we extracted the local maxima in each frame and selected the corresponding regions-of-interest (ROIs) of the signal (Grull et al., 2011; Prakash, 2017). The centre of each signal is precisely determined by a statistical fit approximately the ideal PSF (Airy-function) with a Gaussian. In cases, where the sample background and camera noise are minimal, the fitted position can be estimated with a precision of 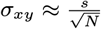, where *s* is the standard deviation of the Gaussian and *N* is the total number of photons detected (Betzig et al., 2006; Thompson et al., 2002). An updated formula by (Mortensen et al., 2010) and (Stallinga and Rieger, 2012) provides a good approximation to calculate the localisation precision for samples with significant background and readout noise.

In our setup, some signals stay ‘on’ for a time longer than the camera integration time. If the signal appears in the consecutive frames, then it can be removed or merged to prevent recounting of the same signal. In the present case, we combined the signals in successive frames if the signals were within the localisation precision of the first signal. For visualisation, each signal position was blurred with a Gaussian distribution of standard deviation equal to the mean distance to the next 20 nearest neighbour molecule positions. The algorithm has been described previously in (Prakash et al., 2015; Kaufmann et al., 2012).

Publicly available SMLM data reconstruction softwares such as ThunderSTORM (Ovesný et al., 2014) or rapidSTORM (Wolter et al., 2012) can be used to carry out the above mentioned analysis.

## Discussion

In this proof-of-principle study, we have shown that high-density superresolution images can be obtained using a Mercury arc lamp and a conventional epifluorescence microscope setup. The system described here is focused on the blinking behaviour of Alexa Fluor 594 in combination with Prolong Diamond. However, the configuration presented should be easily extended to other fluorophores and light sources such as LEDs or metal Halide lamps, given that they can provide with a minimum threshold energy within the excitation bandwidth of the fluorophore (Lichtman and Conchello, 2005). To note, the metal Halide lamps provide a better alternative to the Mercury arc lamps as they have a brighter intensity between the 460-520 nm range, a more uniform field of illumination, and allow for more control of the lamp power (Figure S3). Moreover, they produce less heat than the Xenon and Mercury arc lamps, and no bulb alignment is required.

Once the illumination is optimised, then the care must be taken that different fluorescent moieties have a low duty cycle in addition to having a high photon yield and photostability in order to obtain high-resolution single molecule images (Dempsey et al., 2011). A limited signal density due to the photobleaching of fluorophores has been a central problem with most single-molecule superresolution microscopy techniques, so far. The prolonged reactivation of Alexa Fluor 594 is particularly helpful to re-use the same sample for multiple measurements and to perform imaging on different setups. The prolonged blinking of a fluorophore can be further utilised for single particle tracking experiments (Manley et al., 2008; Gahlmann and Moerner, 2014; Balzarotti et al., 2017). The reactivation with UV illumination is similar to that of Alexa Fluor 647 in a redox buffer but using light instead of chemicals (Heilemann et al., 2008; Vogelsang et al., 2010). It might be possible that lasers bleach the fluorochrome permanently and faster, which might not be the reason in the case of an incoherent light source. A proper calibration for different fluorophores with both lamps and lasers is suggested.

We were also able to perform STED and LFSM on the same biological sample with comparable resolution, confirming the high-resolution setting of the setup. The prolongated blinking with deep blue illumination (350-380 nm) was the key to achieving a high signal density and subsequently the high resolution. This comes from the fact that the final resolution of an image depends not only on the precision of localisation but also on the total number of independently localised signals (Legant et al., 2016; Prakash, 2017).

At this point, we wish to re-emphasize that the blinking phenomenon can be observed in ProLong Diamond as the imaging medium. This simplifies the need to prepare special buffers consisting of the classical redox cocktail required for dSTORM (Heilemann et al., 2008) and other related superresolution techniques (Szczurek et al., 2014). As the composition of Prolong Diamond is not known, at this stage of the study it was difficult to narrow down the minimal requirements for blinking to occur. However, these results hint at a conformational explanation (more likely in a viscous/liquid medium) of the blinking events (Baddeley et al., 2009; Estévez-Torres et al., 2009), in addition to cycles of *H*^+^ binding and release, as previously hypothesized (Fölling et al., 2008; Zurek-Biesiada et al., 2015).

The two main challenges associated with LFSM and SMLM techniques, in general, are the stage/sample drift and optical sectioning (Betzig et al., 2006; Juette et al., 2008). Due to sample drift (correction only up to 50 nm) and the high duty cycle of Alexa Fluor 594, we could not resolve the individual components of the NPCs, which are only 20-40 nm apart. In the case of LFSM, the majority of drift stemmed from heat generated by the Mercury arc lamp and the stage movement. In our case, stabilizing the whole microscope for at least one hour before the measurement significantly helped in minimizing the drift. For good optical sectioning, we made use of very thin biological samples. For thick samples, high NA objectives can be used to achieve a better signal-to-noise ratio at the basal plane. Furthermore, the camera readout noise (CCD vs EMCCD) can be a significant factor in case of the fluorophores with a poor photon count.

The overall cost of the setup is low and comes with a minimal effort in terms of implementation. Most of the other high-resolution imaging techniques require the use of relatively high power lasers, expensive objective lenses (high-NA), very accurate piezoelectric stages and high-end cameras. Lasers are difficult to align, and misalignments often produces artefacts that can be difficult to recognise if the prior information about the sample is not available (Prakash, 2017). Moreover, the technical aspect of the implementation procedures makes these setups a difficult access to many biologists. We hope that LFSM will pave the way for a simple and low-cost high-resolution microscopy implementation especially in emerging countries with limited budgets for science.

We think that the accessibility of the method and the relative ease of its use can democratise superresolution imaging and make it an everyday technique for use in molecular biology studies. Last but not the least, the various photophysical observation such as indefinite blinking due to the photoreactivation and non-permanent bleaching of fluorophores with deep blue illumination indicates for a more thorough investigation on the photophysics of a fluorophore (Lippincott-Schwartz et al., 2003; Lippincott-Schwartz and Patterson, 2008) and various mechanisms that can make a fluorophore blink (Vogelsang et al., 2010).

## Acknowledgements

This work was supported by the National Institute of General Medical Sciences of the National Institutes of Health (grant number GM33397). The content is solely the responsibility of the authors and does not necessarily represent the official views of the National Institutes of Health. K.P. acknowledges Joseph G. Gall, Johannes Hohlbein, David Fournier, Zehra Nizami, Safia Malki, Yixian Zheng and Mahmud Siddiqi for the biological samples, help with microscopy measurements and useful discussion. The power meter (X-Cite XR2100) was provided by Greg Kilduff.

## Conflict of interest

The author declare no conflict of interest.

## Supplemental Material

### Laser-free super-resolution microscopy

**Figure S1.**
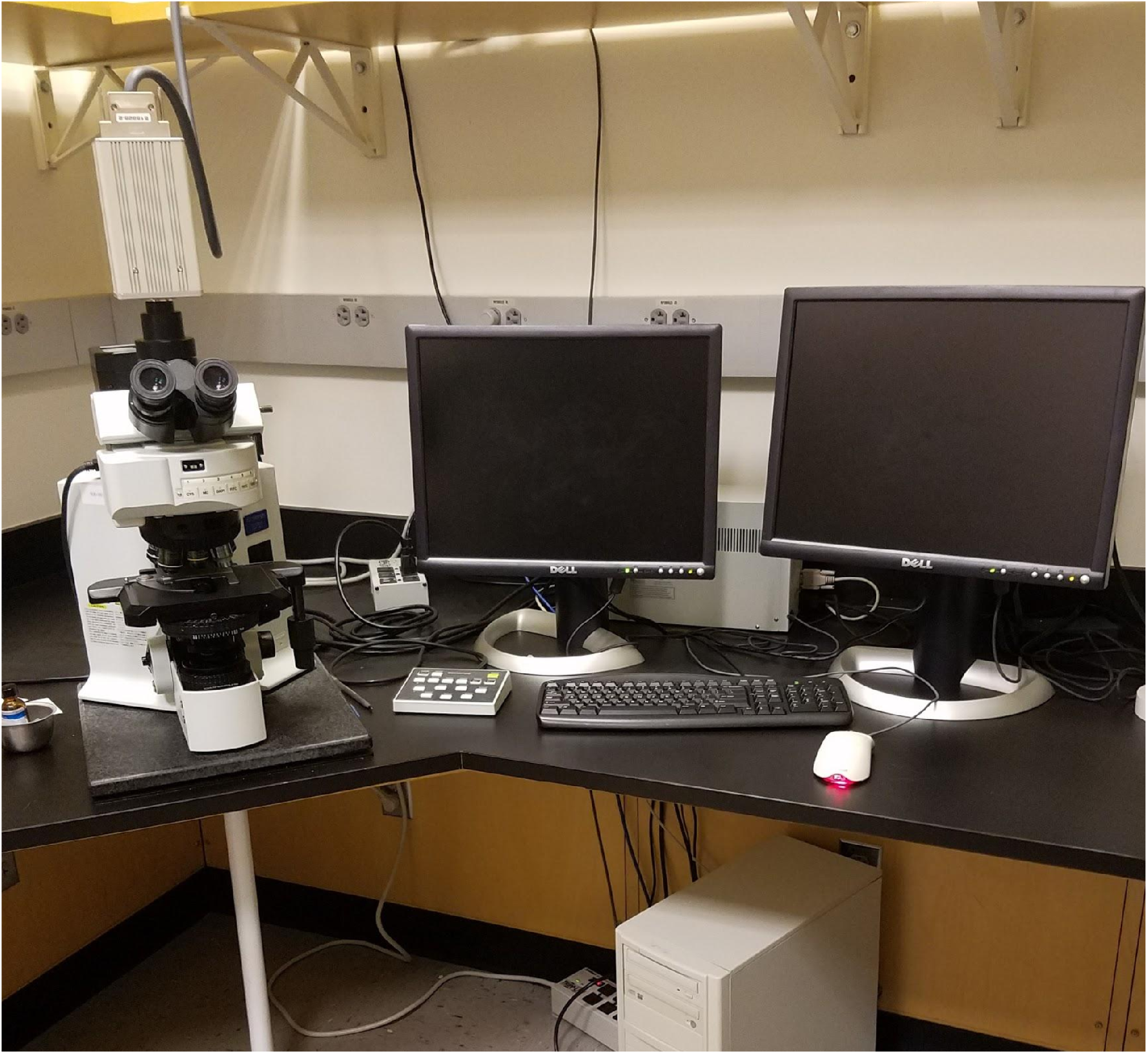
Olympus BX 61 as used for LFSM measurements.

**Figure S2.**
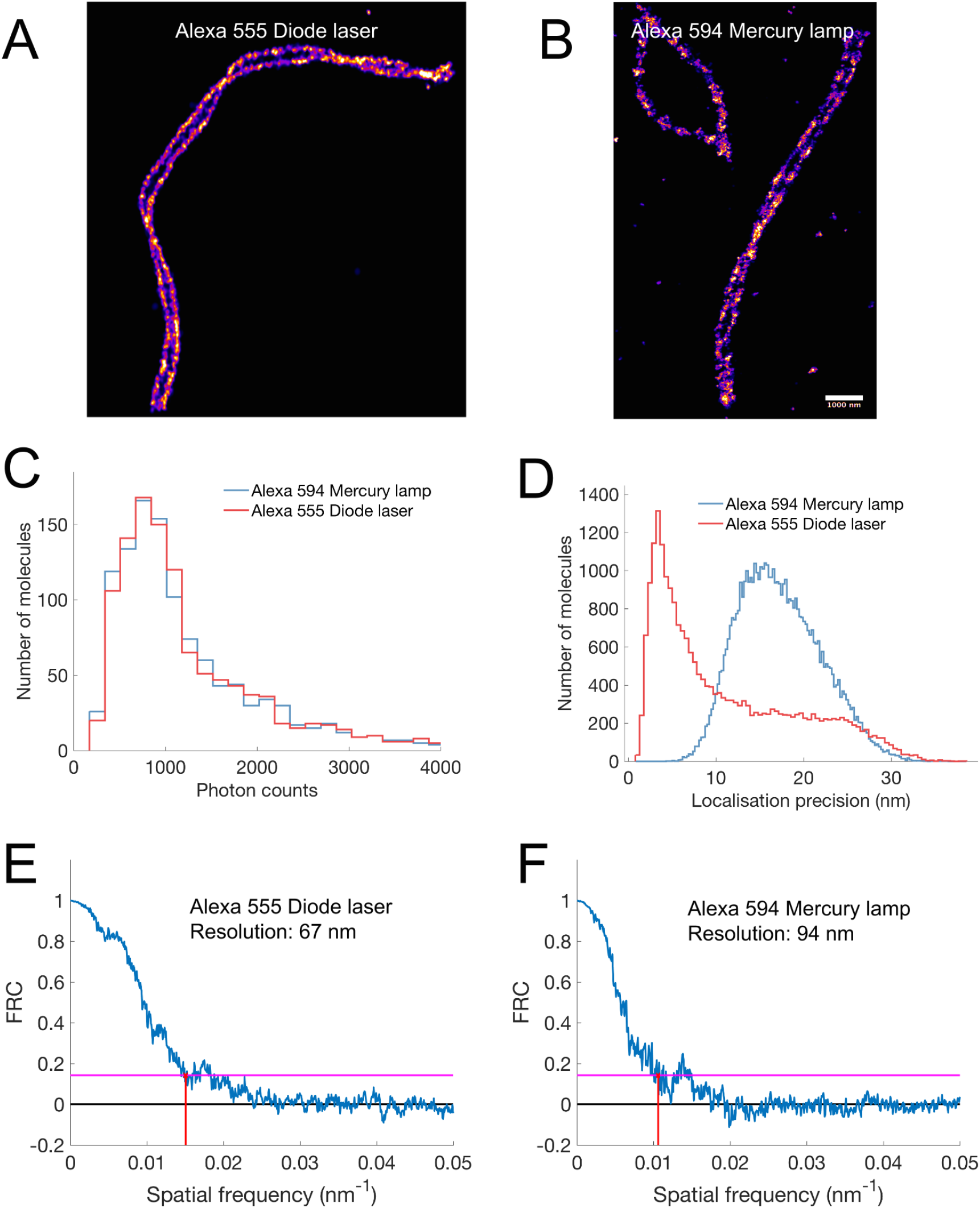
Comparision of laser-based SMLM and lamp-based LFSM: (A) SMLM image of SYCP3 labelled with Alexa Fluor 555 (Prakash et al., 2015). Image acquired with a high-end wide-field setup consisting of a diode laser (561 nm), a very sensitive camera, piezo stage and a high NA objective. (B) LFSM image of SYCP3 labelled with Alexa Fluor 555. The image is acquired with Olympus BX61. The setup consists of an incoherent light source, a simple CCD camera with an ordinary stage. (C) Comparison of photons emitted by single molecules using these two setups. (D) Comparison of the two configurations using localisation precision. (E-F) FRC resolution for Alexa Fluor 555 (SMLM) and Alexa Fluor 594 (LFSM).

**Figure S3.**
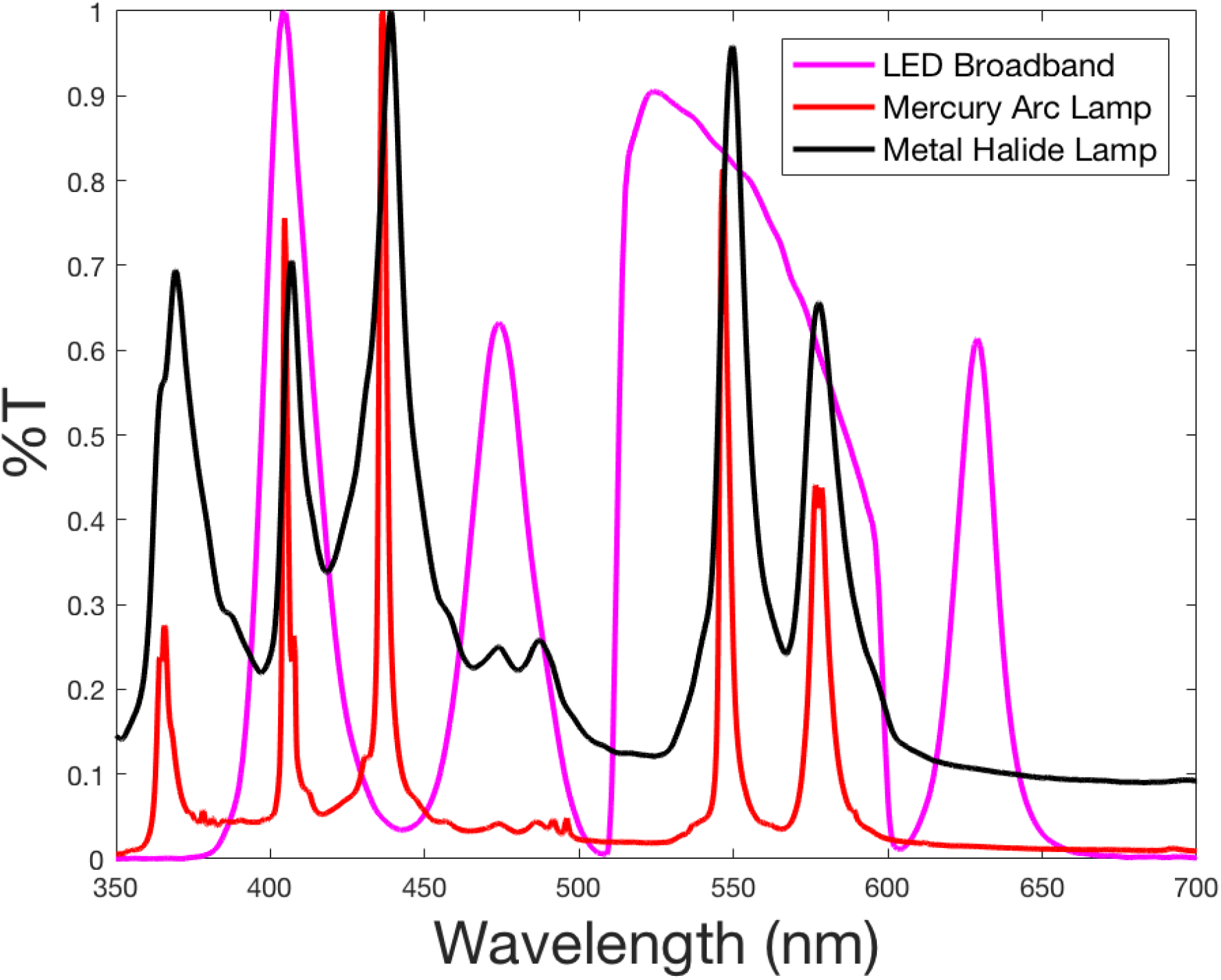
A comparison of spectra of a standard LED broadband, Mercury arc lamp and metal Halide lamp. Both LEDs and metal Halide lamps provide more uniform illumination than the Mercury arc lamps and can be good alternatives.

**Figure S4.**
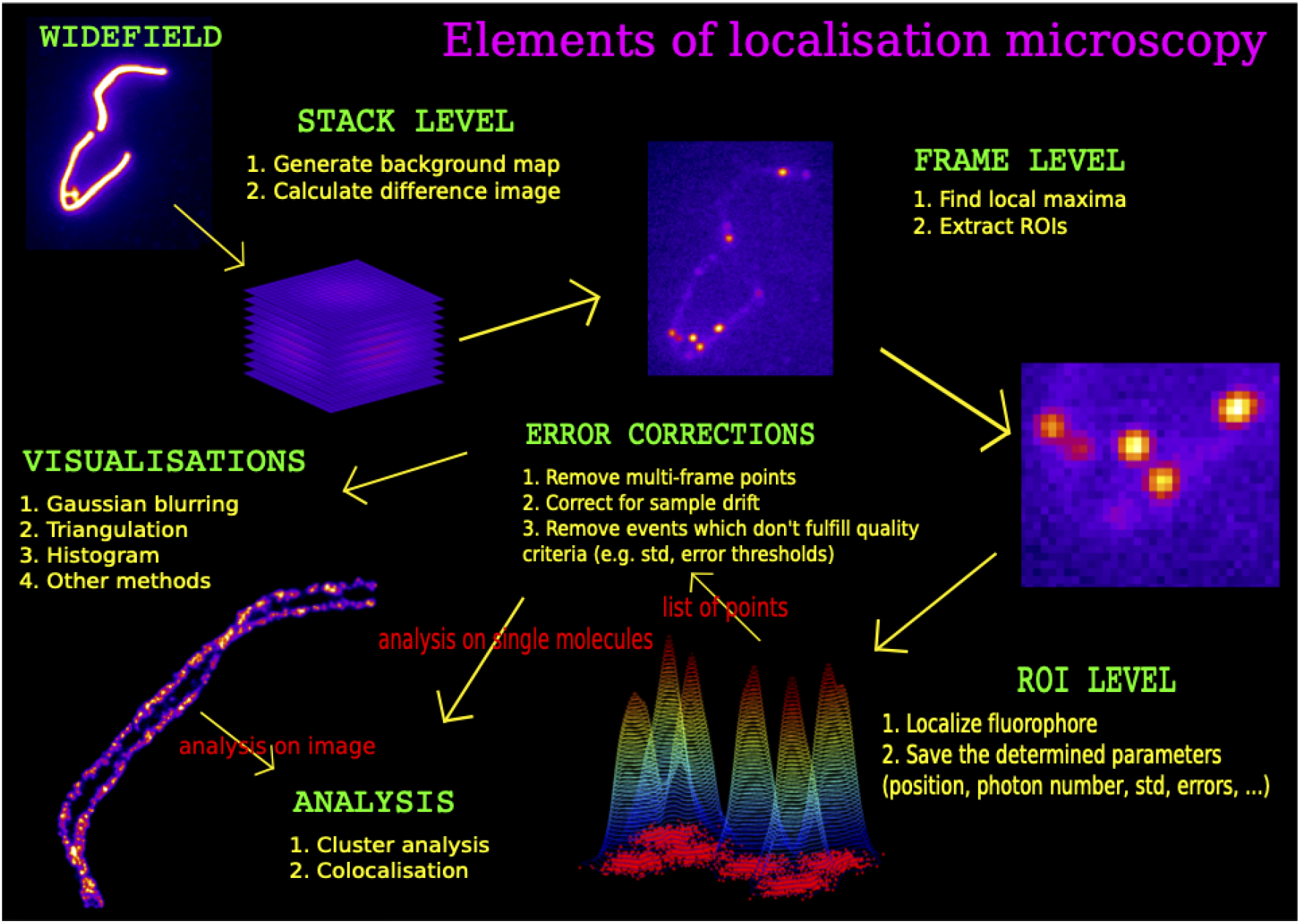
LFSM data reconstruction and analysis flowchart (Prakash, 2016). See Methods and Material section for details.

## Notes

### Competing Interest Statement

The authors have declared no competing interest.

### Summary of Updates

1. Text edits 2. Figure updates

